# Unravelling the Role of Uterine-Derived Estrogen in Establishing Endometrial Receptivity and Supporting Embryo Implantation and Decidualization in the Golden Hamster (*Mesocricetus auratus*)

**DOI:** 10.64898/2026.11.18.767498

**Authors:** Randhir Kumar, Chandana Haldar, Pranab Lal Pakrasi

## Abstract

The mammalian embryo implantation is generally regulated by ovarian estrogen and progesterone in laboratory animals so far studied. In some animals, like golden hamsters, the process of implantation is reported to be independent of ovarian estrogen. Other studies, however, suggest that estrogen-related events do exist for implantation. We also observed the complete absence of implantation in letrozole-treated uterine horns (an inhibitor of the key enzyme for estrogen biosynthesis). However, the vehicle-treated horn bear has a normal implantation site in D6 of pregnancy. Hence, there must be an involvement of estrogen that may come from a source other than the ovary. It may be either non-ovarian or locally synthesized (non-systemic, uterine). But the involvement of non-systemic estrogen is not well documented. So, in the present study, the de novo synthesis of estrogen in the uterus and its role in early pregnancy is described. For this, letrozole-treated and control uterine horns were further studied using the tools of proteomics and genomics to know the exact cause of failure of implantation. Using IHC, it was observed that there is a presence of cytochrome P450 aromatase, a rate-limiting enzyme in the estrogen biosynthesis pathway, in the uterine horn of pregnant hamsters at the time of implantation. The expression profile of this enzyme and their mRNA was further checked by western blot and real-time PCR, respectively. It also corroborated earlier IHC observations. We found that their expression is markedly induced at the implantation site and in stromal cells around the day of implantation and decidualization, respectively. This aromatase is responsible for de novo synthesis of estrogen in association with other steroidogenic enzymes, which was also observed to be present in the uterus during early pregnancy. This locally synthesized estrogen is sufficient to elicit other implantation-specific physiological factors required for successful embryo implantation and decidualization in the absence of ovarian estrogen. Since embryo implantation in human beings is also ovarian estrogen independent and much more similar to hamster, so, this study may help in managing issue concerning to infertility in women due to early loss of pregnancy.

## Introduction

Infertility is one of the devastating problems worldwide in recent decades which adversely affect the socioeconomic status of a person. Loss of early pregnancy due to failure in embryo implantation is considered as one of the major cause of infertility in females and most important limiting factor for achieving a successful outcome using the assisted reproductive techniques (ART) (Noewitz et al., 2001 and Zhang et al., 2013). Statistic report suggests that one among each nine couple in Europe and USA is affected by early pregnancy disorders related to implantation failure (Krussel et al., 2003). Embryo implantation is one of the most crucial step in the female reproduction in which the competent blastocyst get adhered to receptive endometrium during a narrow window of time to established successful pregnancy (Ma et al., 2003). This requires a cross talk between blastocyst and endometrium to achieve a synchronized growth and development of blastocyst and attaining a stage of endometrial receptivity for successful implantation. In response of implanting blastocyst, uterus undergo extensive remodeling in which stromal cells surrounding the implanting blastocyst proliferate and differentiate in to a specialized cell types commonly known as decidual cells by a process of decidualization (Cha et al., 2012). Several implantation specific molecules like vasoactive mediators, growth factors, cytokines and transcription factors are recruited by either endometrium or/and blastocyst at feto-maternal interface during window of implantation and play very critical role in determining endometrial receptivity (Dey et al., 2004; Simon et al., 2000 and Singh et al., 2011). Ovarian steroids are responsible for the inducing these factors at the implantation site during early pregnancy (Carson et al., 2000 and Lim et al., 2002). On the basis of hormonal level in estrous cycle and menstruation cycle, it was assumed that progesterone is the principal hormone for the process of embryo implantation in almost all mammals. Estrogen secreted in proliferative phase is responsible for the proliferation of endometrial cells while luteal phase estrogen showed species specific role in this process (Daya et al., 2009 and Wang et al., 2006). But it has been established that only progesterone is sufficient to elicit the cascade of embryo implantation in golden hamster, rhesus monkey, rabbit as well as in human. In other hand, administration of estrogen receptor antagonist prior to embryo implantation adversely affects this process.

After embryo implantation, progesterone secreted from corpus luteum is responsible for the process of decidualization and maintenance of pregnancy (Wang et al., 2004). An early report showed that progesterone alone is sufficient to induce artificial decidualization in ovariectomized mice (Paria et al., 1999). In contrast, administration of ICI-182780 in mice adversely affect the process of decidualization (Curtis et al., 1990), suggesting a role of estrogen via estrogen receptors in the process of decidualization. These apparently contradictory reports clearly suggest a critical role of estrogen which is produced from other source rather than ovary. Although, ample amount of study has been performed to understand the complex events of embryo implantation in different species, none of them are able to explain the ambiguous role of estrogen in the embryo implantation in golden hamster and also in human. Thus, in present study we have considered a novel approach of *de novo* synthesis of estrogen in uterus to explain the probable role of non-systemic estrogen in the embryo implantation of golden hamster. Since, the embryo implantation in human being is similar to that of golden hamster in respect of requirement of ovarian estrogen, the outcome of this study may be translated directly in human being to manage the idiopathic problem of infertility and for the improvement of success rate of IVF.

## Materials and Methods

### Animals care and maintenance

Golden hamsters were kept in a controlled environment at 14 hrs light and 10 hrs dark cycles in an animal house facility of Department of Zoology, Banaras Hindu University, Varanasi, India. They were provided unlimited access of food and water. 10-12 week old hamsters were considered for mating. All experimental procedure were conducted according to the institutional guideline and strictly followed the rules of experimental animals (Scientific Procedure) Act 2007, of the Committee for the purpose of Supervision and Control on experiments on Animals (Reg. No.-802/Go/Re/S/IS/CPSCEA), Government of India, on animal welfare.

### Sample collection

Adult female hamsters with four consecutive estrous cycles were kept with proven male on the evening of proestrus cycle in 1:1 ratio to get dated pregnancy. Confirmation of sperm in vaginal smear in the next morning was considered as day 1 (D1) of pregnancy. Whole uterus were collected from day1to day 3 of pregnancy and embryo were flushed from the end of oviduct using normal saline. Implantation and inter-implantation site were separately excised from the uterus of day 4 to day 8 pregnancies. Tissues were immediately stored at –80°C for real time PCR and Western blot analyses. Some parts of tissues were fixed in bouins’s fluid for immunohistochemistry. Every experiment was performed in minimum five independent biological and technical replicates.

### Letrozole treatment

Letrozole (Abcam) is a specific inhibitor of cytochrome P_450_ aromatase, a rate limiting enzyme for the biosynthesis of estrogen. This inhibitor was used to cut the supply of any locally synthesized estrogen in uterus which might be regulate the activation of implantation specific molecules used in this study. Inhibitor was administered intrauterine (20mg/kg) in one uterine horn in D3 of pregnancy while contralateral horn was treated with vehicle and used as a control for the comparison of expression of different molecules used in this study. Another group of animals were treated with same dose of letrozole along with estrogen (Sigma-Aldrich, 1µg). Hamsters were sacrificed at D6 of pregnancy and both horns were harvested for further analysis. Five independent animals were selected for each group for this study.

### Immunohistochemistry

Immunohistochemical staining was performed following method of Pakrasi and Jain, 2008 with slight modification. Briefly, the whole uteri/implantation and inter-implantation sites were removed and fixed in Bouin’s solution overnight. Further, tissues were dehydrated in ascending grades of ethanol and cleared in xylene and embedded in paraffin wax. A 5 to7 μm tissue sections were mounted on poly-L-lysine coated slides. After deparaffinization in xylene and rehydration in descending grades of ethanol, tissues were subjected for heat induced antigen retrieval for 15 min in microwave oven using 10mm sodium citrate buffer (pH 6.0). The slides were allowed to cool at room temperature and then incubated with normal horse serum for 2 h to block any nonspecific reaction. The sections were incubated with respective primary antibody (Table no. 1 for antibody details) for overnight at 4°C. This is followed by incubation with appropriate biotinylated secondary antibody (Abcam, UK) for 1 hr at room temperature. Endogenous peroxidase activity was blocked by incubating sections in 3% hydrogen peroxide (H2O2, Sigma, USA) in methanol for 2 min at room temperature. Slides were washed thoroughly in PBS (pH=7.4) between incubation. Finally, sections were stained using ABC kit (Vector laboratory, USA) according to the manufacturer’s instruction. The signal reaction was performed by DAB (Sigma, USA) followed by counterstaining with hematoxylin for cytoplasmic proteins before mounting. Brown deposits indicate the positive signals. Images were captured by Leica DMIRB, Leica Microsystems Wetzlar, GmbH, Germany having external camera of canon (Japan).

**Table 1.**
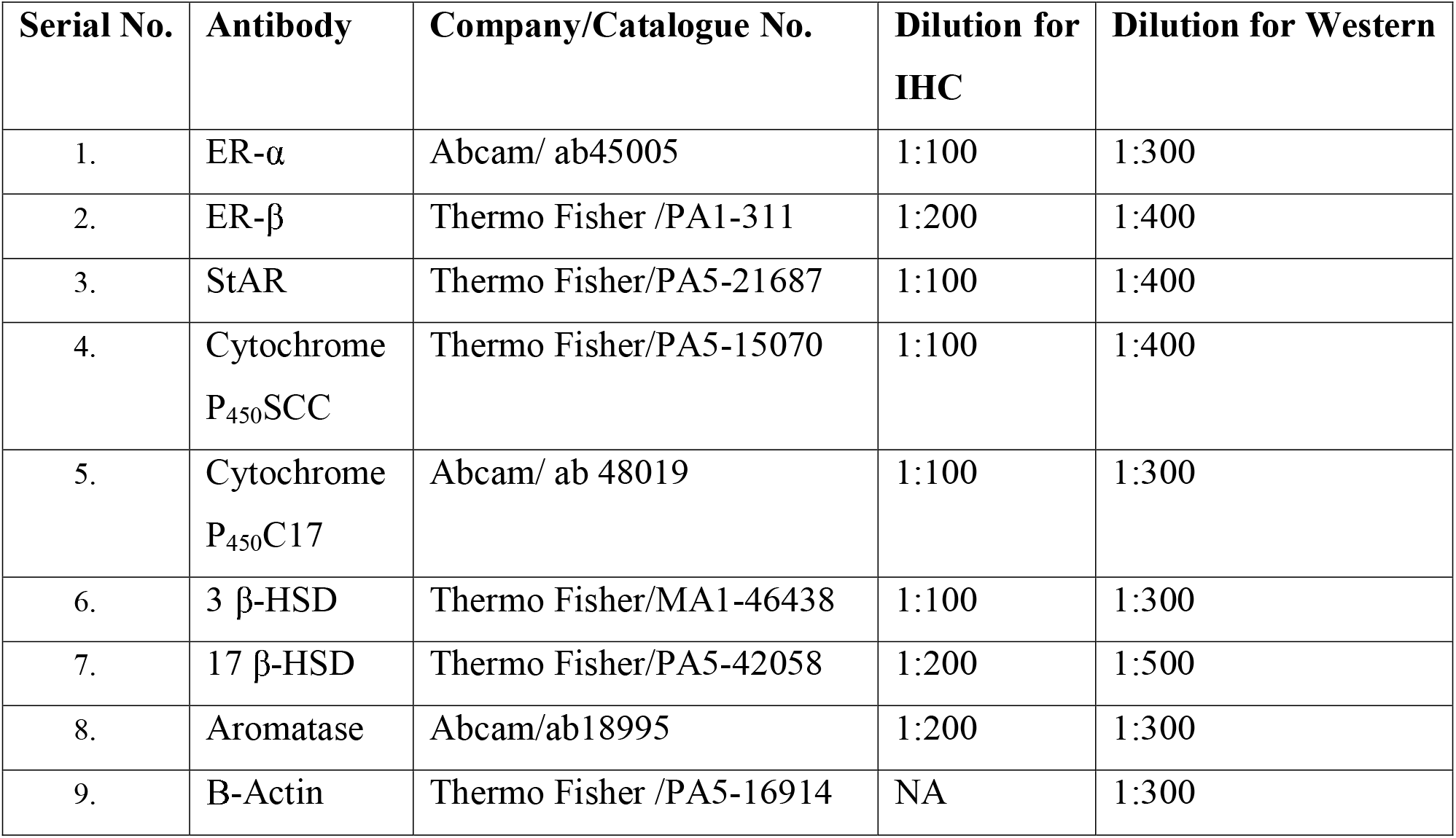
Details of Antibody used in IHC and Western blot.

### Western blot analyses

Western blot analyses were performed following method of Pakrasi and Jain, 2008 with slight modification. In brief, whole uteri were taken up to D4 of pregnancy for extraction of protein. D5 to D7 uteri were repeatedly flushed with normal saline to remove any extent of embryo from the implantation site. After that, implantation sites and inter-implantation sites were separated with the help blue colour impression of pontamine dye (Sigma, USA). Different tissues were homogenized in lysis buffer containing 150 Mm NaCl, 50 Mm Tris (pH 8), 1% NP-40, 0.1% SDS, 0.5% Sodium deoxycholate and fresh protease inhibitor (1 μg/ml aprotinin, 1 mM EDTA, 0.5 μg/ml leupeptin, 100 μg/ml PMSF and 1 mM sodium orthovanadate). Tissue homogenate were kept on rocker at 4°C for 30 min. Homogenate were centrifuged at 10,000 ×g at 4°C for 10 min to remove cell debris and supernatant were collected and stored at 80°C. Protein concentration was determined using Quick Start Bradford Protein Assay (Bio-Rad, USA). After normalizing for equal protein concentration, cell lysates were boiled in Laemmli 2×sample buffer (4% SDS, 20% Glycerol, 125 mM Tris (pH 6.8) 0.02% Bromophenol blue and 10% β-mercaptoethanol) for 5 min. Proteins were subjected for 10% SDS-PAGE and transferred to PVDF membrane (polyvinylidene difluoride, Millipore, USA). The membranes were blocked in 5% non-fat milk in TBS-T (10 mM Tris-HCl (pH 8), 150 mM NaCl and 0.05% tween 20) for 2 h at room temperature. Membranes were incubated with diverse primary antibodies with different dilutions (see Table no. 1 for antibody details) for overnight at 4°C. After incubation, membranes were washed two times (10 min each) with TBS-T and subjected for corresponding HRP-conjugate secondary antibody incubation in 5% milk for 2 h at room temperature. The membranes were washed and incubated with ECL reagent (Millipore, USA) and exposed to X-ray film for the detection of desired bands. Equal loading of protein was confirmed by β-actin (Thermo-Scientific, USA). Density of bands was quantified using image-J software, NIH.

### Gene expression analysis

Total RNA was extracted from the uterine horn of pregnant golden hamster using Trizol-reagent (Invitrogen, USA). RNA was treated with DNase (Ambion, USA) to avoid any DNA contamination. RNA was run on 2% gel to check their integrity and quantified by nanodrop spectrophotometer (Nanodrop Technologies,). RNA samples with 260/280 ratio ranges 1.9 2.1 was taken for further experiment. The cDNA was synthesized following manufacturer’s protocol by using commercial c-DNA synthesis kit (Bio-Rad, USA).

We performed quantitative real time PCR analysis and reverse transcription PCR for all of the genes used in study along with β-Actin as control. Primers were designed using IDT-Primer Quest with default parameters. The primer sequences are listed in Table 2. Specificity of the primers was analyzed using BLAST tool of NCBI. Real time was conducted on 96 well plate and performed on iQ5 thermo cycler (Bio-Rad, Hercules, CA, USA) with iQ5 Optical System Software version 2.0 (Bio-Rad, Hercules, CA, USA). Each reaction mixture of 20 μl contained 5 μl first strand cDNA (100 ng) as template, 0.8 μl of each gene specific primer (200 nM) and 10 μl SYBR Green qPCR Mix (Bio-Rad, USA). Quantitative RT-PCR was performed with the initial incubation for 1 min at 95°C, followed by 40 cycles at 95°C for 15 s and extension at 60°C for 60 s. The dissociation curves were achieved for the assurance of absence of nonspecific amplifications. Three biological and three technical replicates were taken for each sample, and standard deviation and standard error were calculated. As an endogenous control, β-Actin was used for the normalization of Ct value obtained and the analysis of relative expression of mRNA was performed using ΔΔCt method (Livak and Schmittgen, 2001).

**Table 2.**
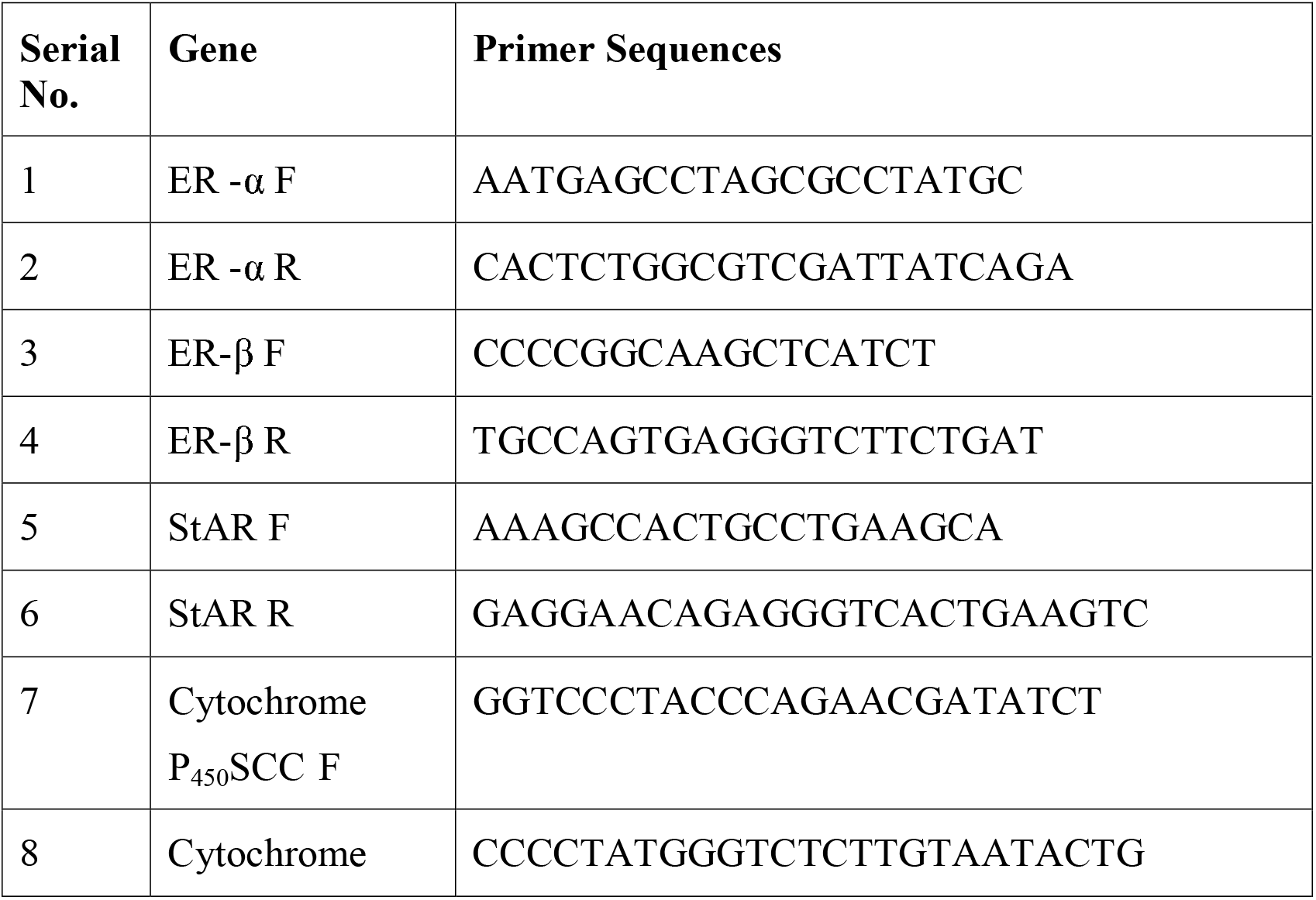
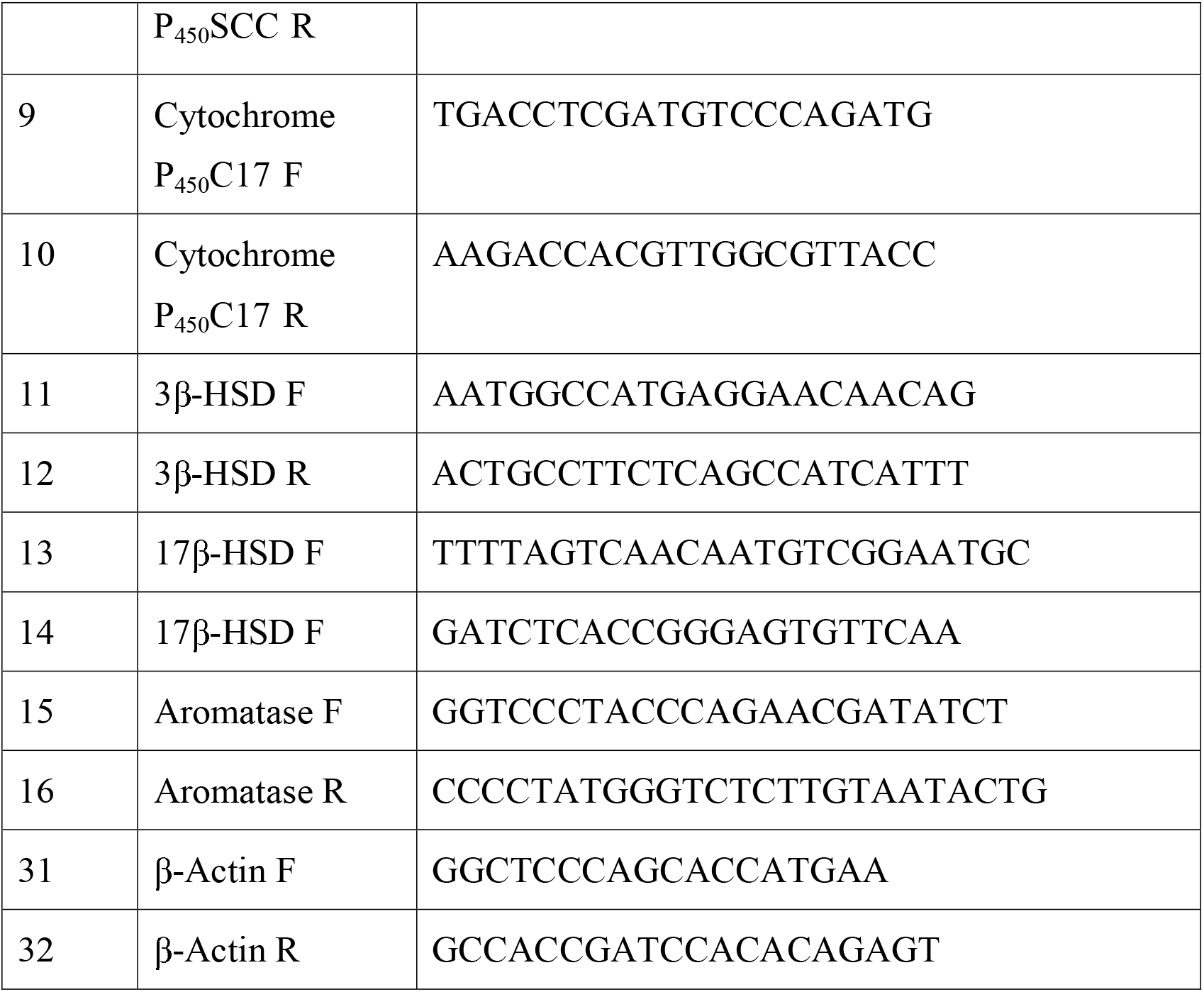
List of primers used for Real-Time PCR.

**Table 3.**
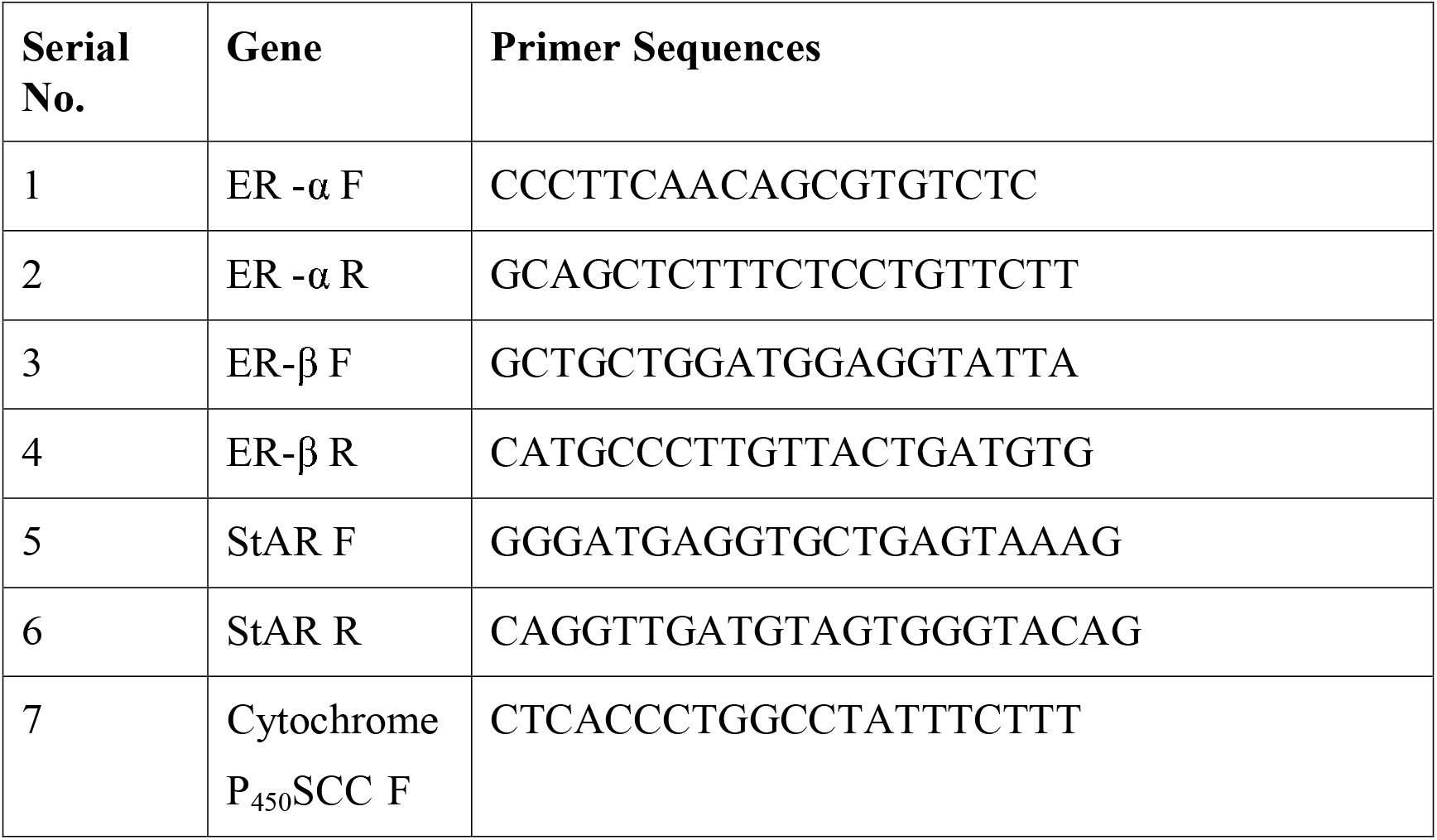
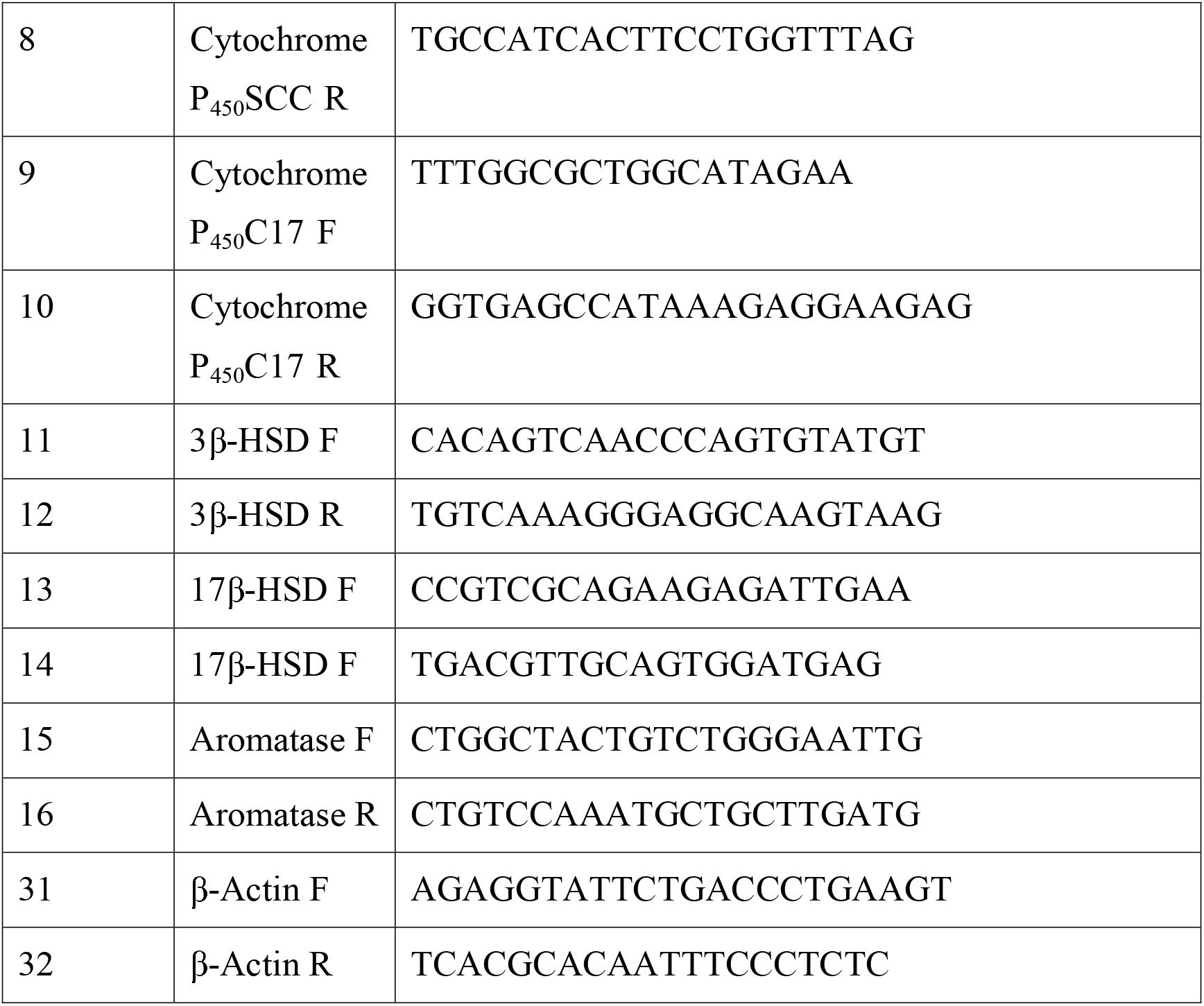
List of primers used for Reverse Transcription PCR.

To analyze the gene expression by reverse transcription PCR, 1 μg of total RNA was subjected to reverse transcription reaction using SuperScript II RNase H-reverse transcriptase (200 units per reaction; Invitrogen, USA) with oligo(dT) primer for 50 min at 42°C. PCR amplification was performed as follows: initial denaturation at 94°C for 2 min followed by 27 cycles of incubations at 94°C for 20 s, 55°C for 40 s, and 72°C for 1.0 min, with a final extension at 72°C for 10 min with iTaq DNA polymerase (iNtRON Biotechnology, Korea) on BioRad C 1000 thermal cycler. β-Actin expression level was used as a quantitative control. Aliquots of individual PCR products were resolved by agarose gel electrophoresis and visualized with ethidium bromide under UV light. All experiments were repeated at least three times, and results from one representative experiment are shown.

### Statistical analysis

At least five independent biological and three technical replicate were taken for each set of experiment. All data were represented as mean ±SEM. Comparison among different groups of pregnancy was analyzed by one way ANOVA followed by Dunnett *post hoc* test taking D1 pregnancy as a control. Data were considered significant when P ≤ 0.05. All statistical analysis were performed using (PRISM software version 5.0; GraphPad, San Diego, CA.

## Results

### Intrauterine administration of letrozole completely inhibit embryo implantation

Letrozole is a well known specific aromatase inhibitor which prevents the biosynthesis of estrogen by inhibiting cytochrome P_450_ aromatase, a key enzyme for the synthesis of estrogen. To assess the role of any uterine estrogen in golden hamster during early pregnancy, letrozole was administered intrauterine in one uterine horn at day3 of pregnancy while contralateral uterine horn was treated with vehicle. We observed the complete absence of embryo implantation in letrozole treated horn while sham control horn showed normal implantation (Fig. 1). Further we observed that this process was reversed when estrogen was administered along with letrozole. This clearly indicates the involvement of local estrogen in the embryo implantation of golden hamster.

**Fig. 1:**
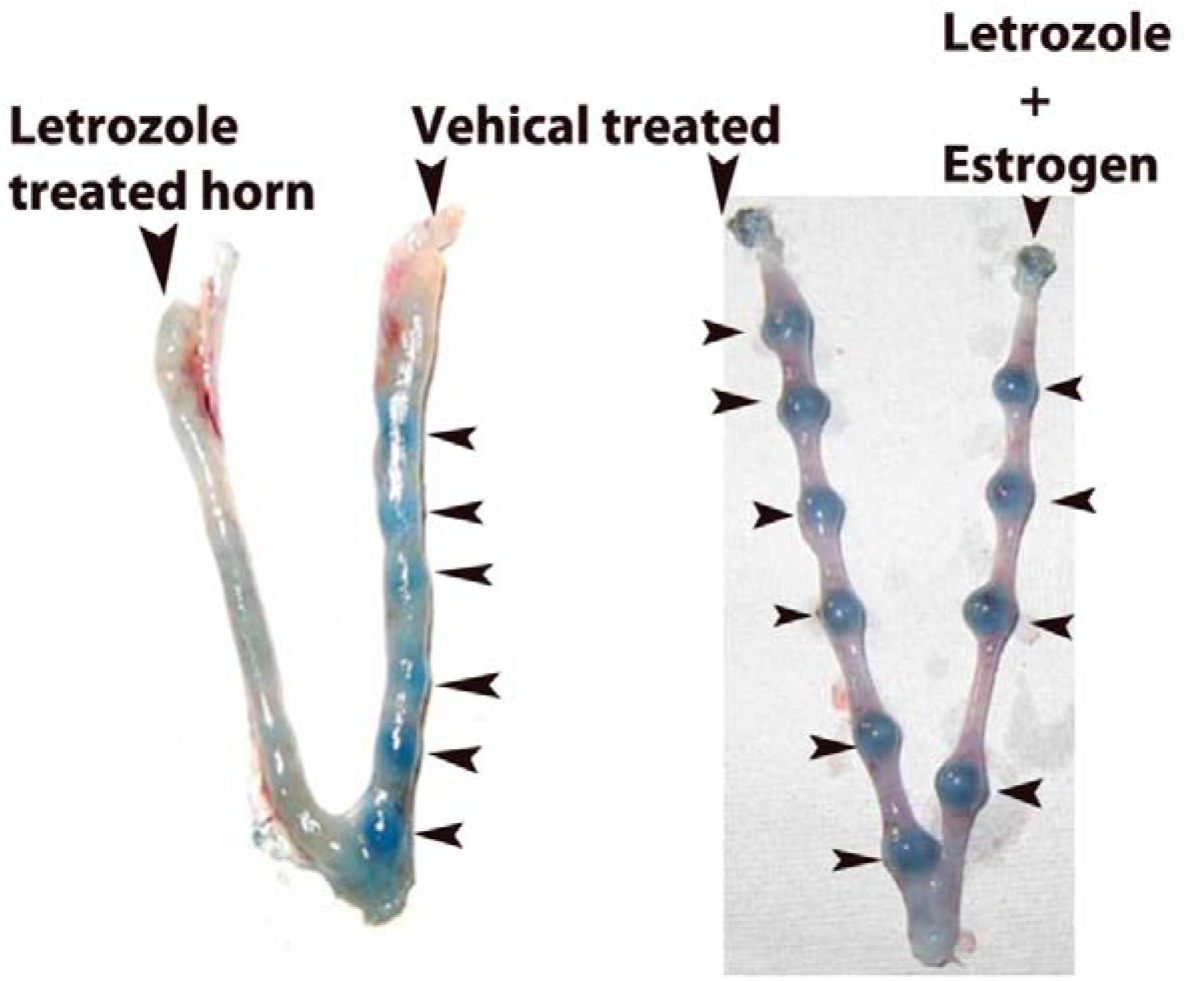
Effect of letrozole on embryo implantation of golden hamster. Embryo implantation was visualized by the injection of pontamine sky blue dye which gives a discrete blue band in sham control and letrozole with estrogen treated horn. However only letrozole treated horn showed complete absence of blue band (Implantation site).

### Spatiotemporal distribution of different enzymes of estrogen biosynthesis pathway in early pregnant uterus

The results of letrozole treatments indicate the possibility of uterine estrogen synthesis during early pregnancy of golden hamster. Therefore, we checked the presence of specific enzymes in the uterus for further confirmation of our hypothesis. We observed the presence of all enzymes of estrogen biosynthesis in spatiotemporal manner in the uterus during early pregnancy (Fig. 2). StAR, first enzyme in estrogen biosynthesis pathway, was found to be present only in myometrial region in the uterus of day1 (D1) and (D2). It appeared in luminal epithelia, glandular epithelia as well as in stromal cell from D3 of pregnancy and persists in almost same cellular location till D4 of pregnancy. In D5 it has more immunopositive cells in antimesometrial side of lumen and primary decidual zone (Fig. 2A).

**Fig. 2A:**
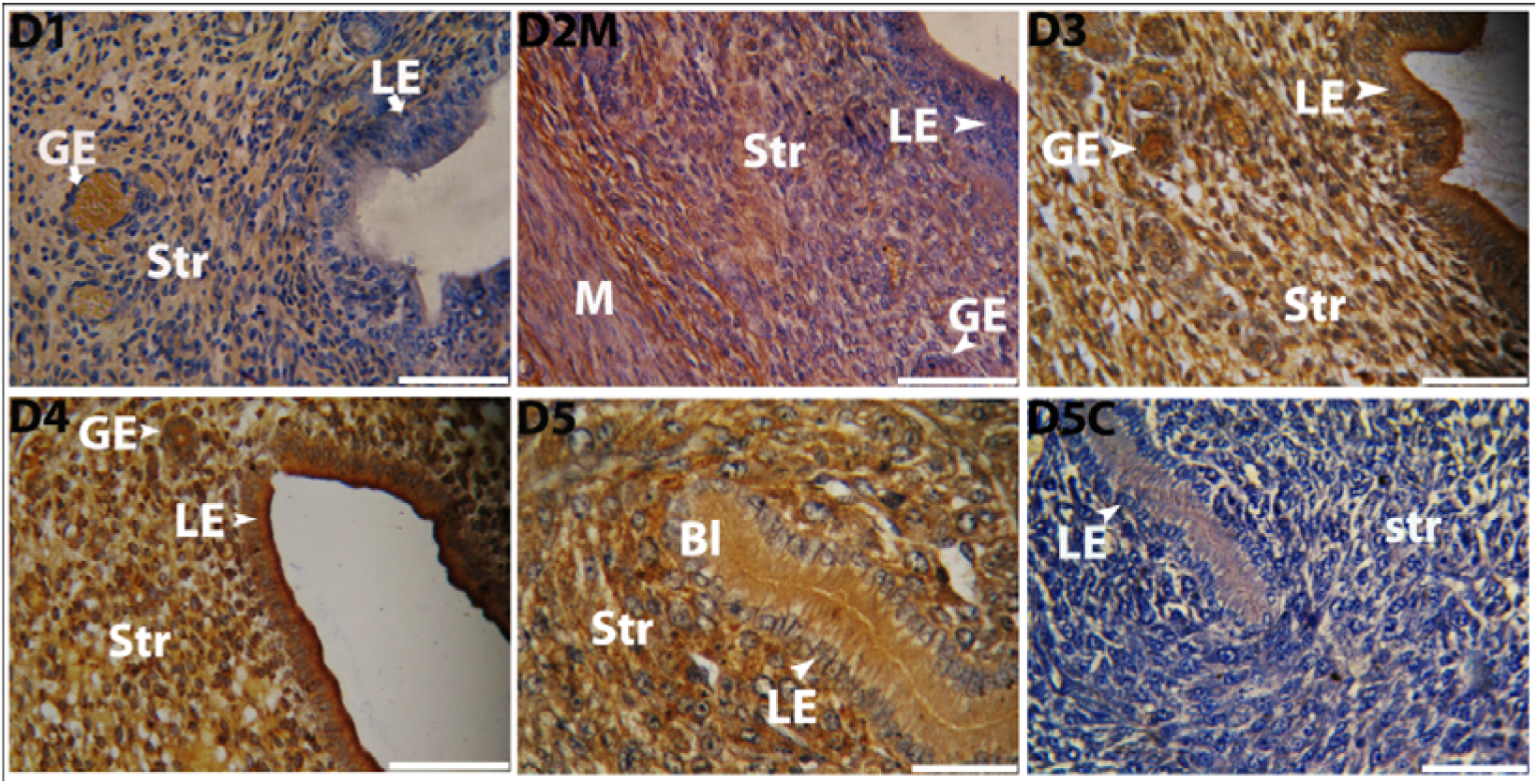
Immunohistochemical localization of StAR protein in the uterus of golden hamster durig pre-implantation (D1 to D3) and peri-implantation (D4 and D5) period showing their spatiotemporal localization in stroma (Str), luminal epithelium (LE) and glandular epithelium (GE). Brown deposition represents positive staining. Primary antibody has been replaced by rabbit IgG in day 5 (D5) pregnant uterus and considered as negative control (D5C). Photographs were taken under bright field condition. Scale bar-50µm and n = 5 in all the groups. Bl represents blastocyst and D represent day of pregnancy. DM showing myometrium region of uterus in specified dated pregnancy and DC represent negative control of specific dated pregnancy.

**Fig. 2B:**
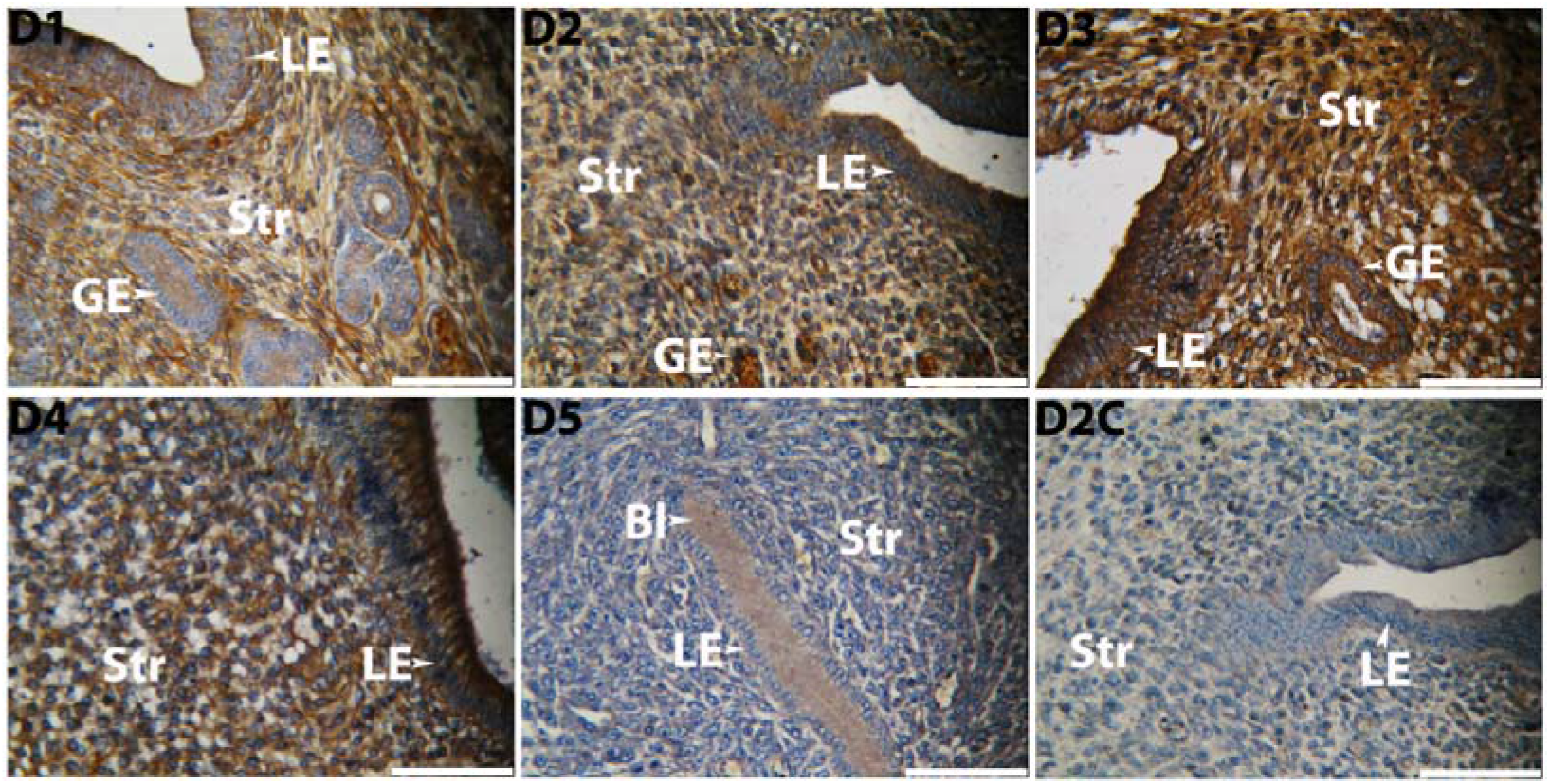
Immunohistochemical localization of cytochrome P_450_ scc protein in the uterus of golden hamster during pre-implantation (D1 to D3) and peri-implantation (D4 and D5) period showing their spatiotemporal localization in stroma (Str), luminal epithelium (LE) and glandular epithelium (GE). Brown deposition represents positive staining. Primary antibody has been replaced by rabbit IgG in day 2 (D2) pregnant uterus and considered as negative control (D2C). Photographs were taken under bright field condition. Scale bar-50µm and n = 5 in all the groups. Bl represents blastocyst and D represent day of pregnancy. DM showing myometrium region of uterus in specified dated pregnancy and DC represent negative control of specific dated pregnancy.

**Fig. 2C:**
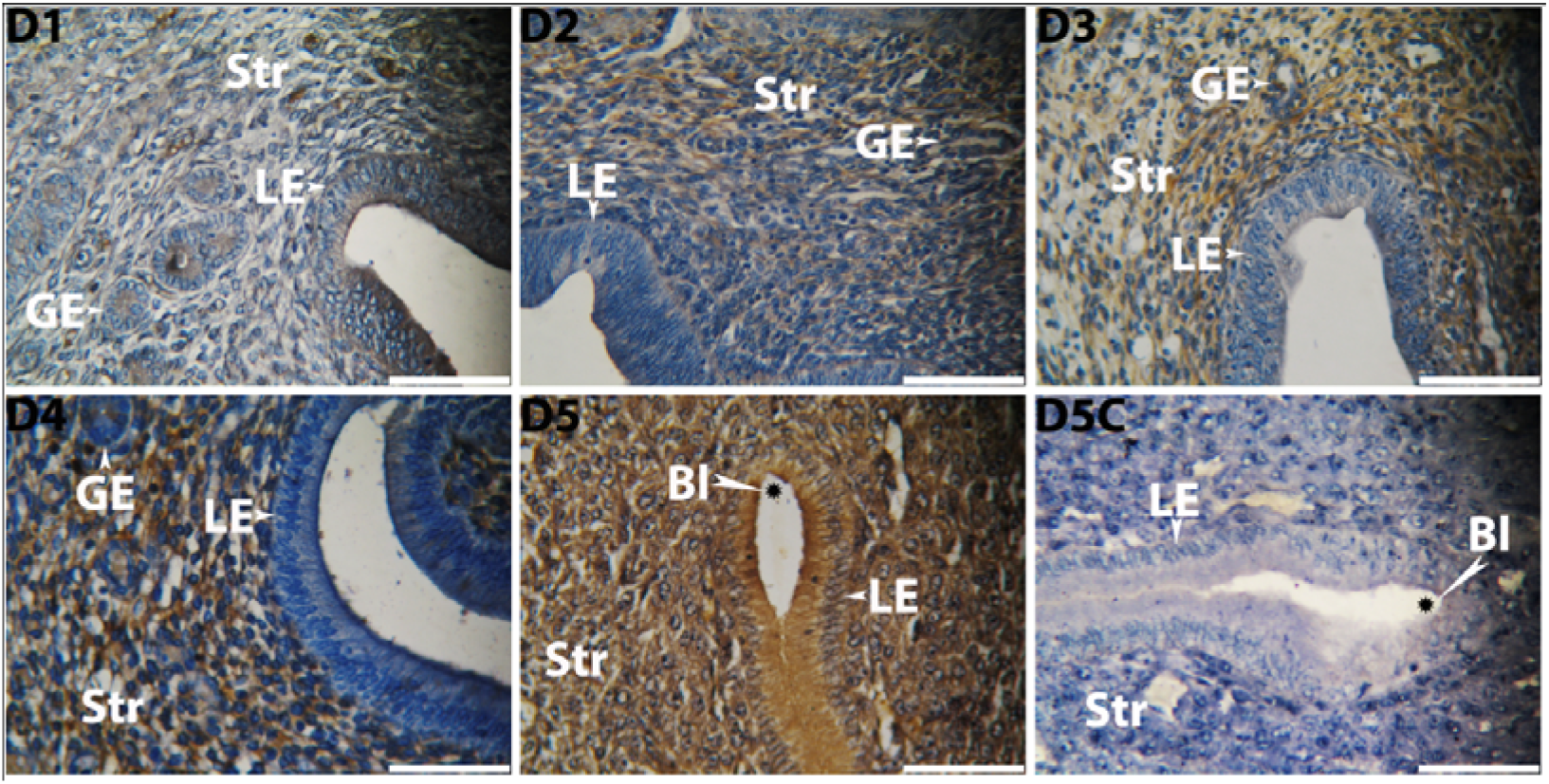
Immunohistochemical localization of cytochrome P_450_ C17 protein in the uterus of golden hamster during pre-implantation (D1 to D3) and peri-implantation (D4 and D5) period showing their spatiotemporal localization in stroma (Str), luminal epithelium (LE) and glandular epithelium (GE). Brown deposition represents positive staining. Primary antibody has been replaced by rabbit IgG in day 5 (D5) pregnant uterus and considered as negative control (D5C). Photographs were taken under bright field condition. Scale bar-50µm and n = 5 in all the groups. Bl represents blastocyst and D represent day of pregnancy. DM showing myometrium region of uterus in specified dated pregnancy and DC represent negative control of specific dated pregnancy

**Fig. 2D:**
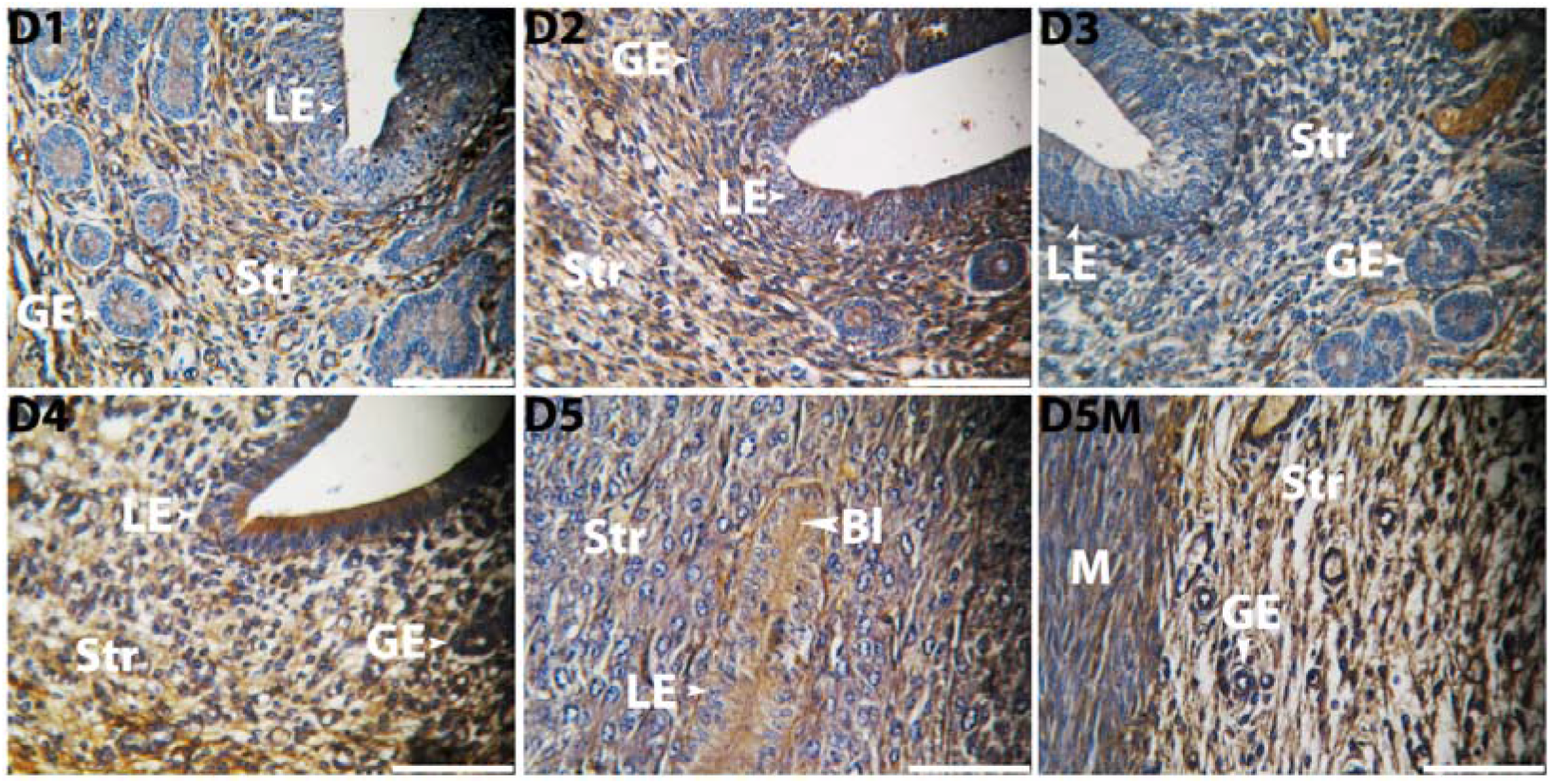
Immunohistochemical localization of 3β-HSD protein in the uterus of golden hamster during pre-implantation (D1 to D3) and peri-implantation (D4 and D5) period showing their spatiotemporal localization in stroma (Str), luminal epithelium (LE) and glandular epithelium (GE). Brown deposition represents positive staining. Photographs were taken under bright field condition. Scale bar-50µm and n = 5 in all the groups. Bl represents blastocyst and D represent day of pregnancy. DM showing myometrium region of uterus in specified dated pregnancy and DC represent negative control of specific dated pregnancy.

**Fig. 2E:**
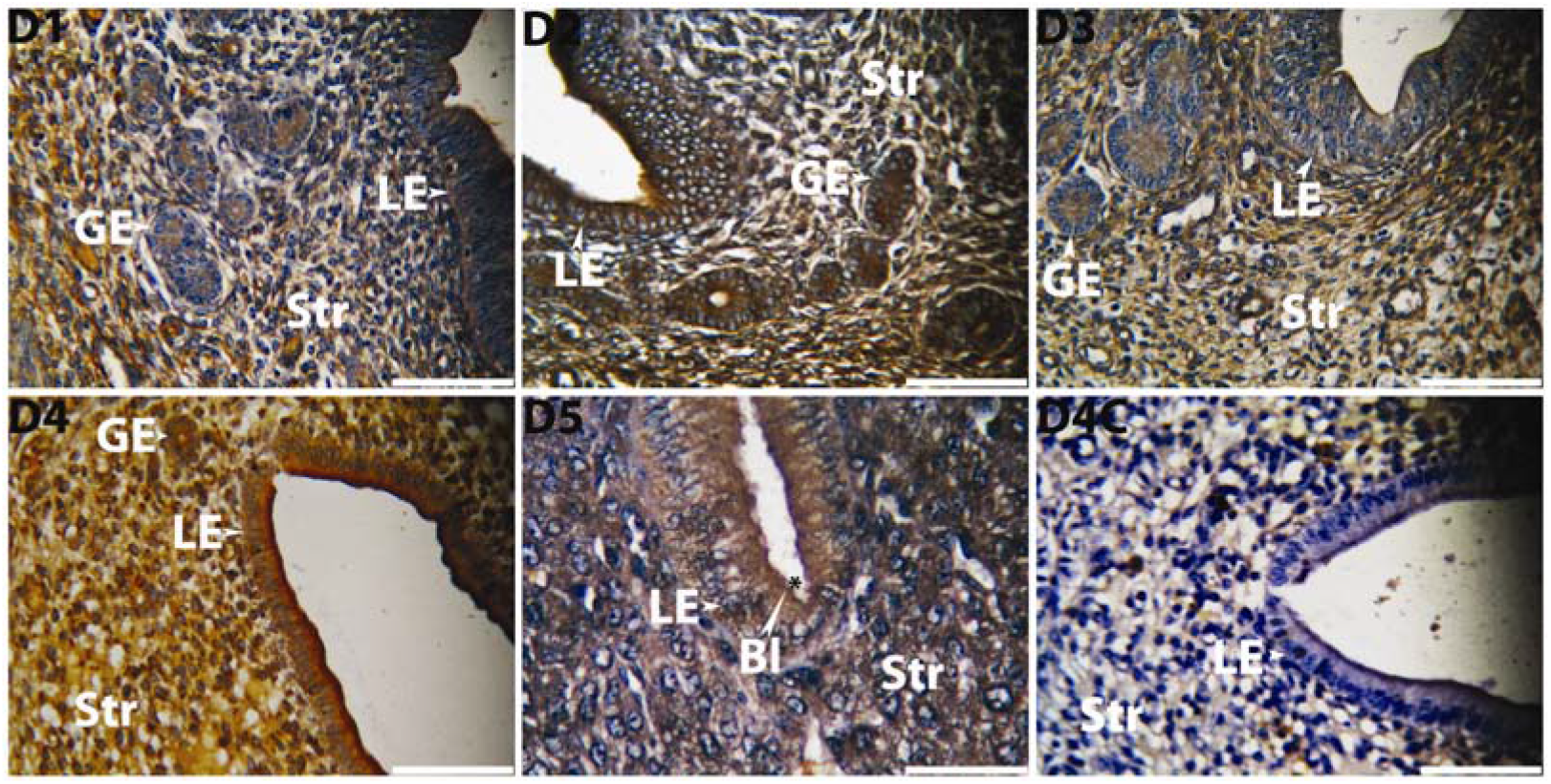
Immunohistochemical localization of 17β-HSD protein in the uterus of golden hamster during pre-implantation (D1 to D3) and peri-implantation (D4 and D5) period showing their spatiotemporal localization in stroma (Str), luminal epithelium (LE) and glandular epithelium (GE). Brown deposition represents positive staining. Primary antibody has been replaced by rabbit IgG in day 4(D4) pregnant uterus and considered as negative control (D4C). Photographs were taken under bright field condition. Scale bar-50µm and n = 5 in all the groups. Bl represents blastocyst and D represent day of pregnancy. DM showing myometrium region of uterus in specified dated pregnancy and DC represent negative control of specific dated pregnancy.

**Fig. 2F:**
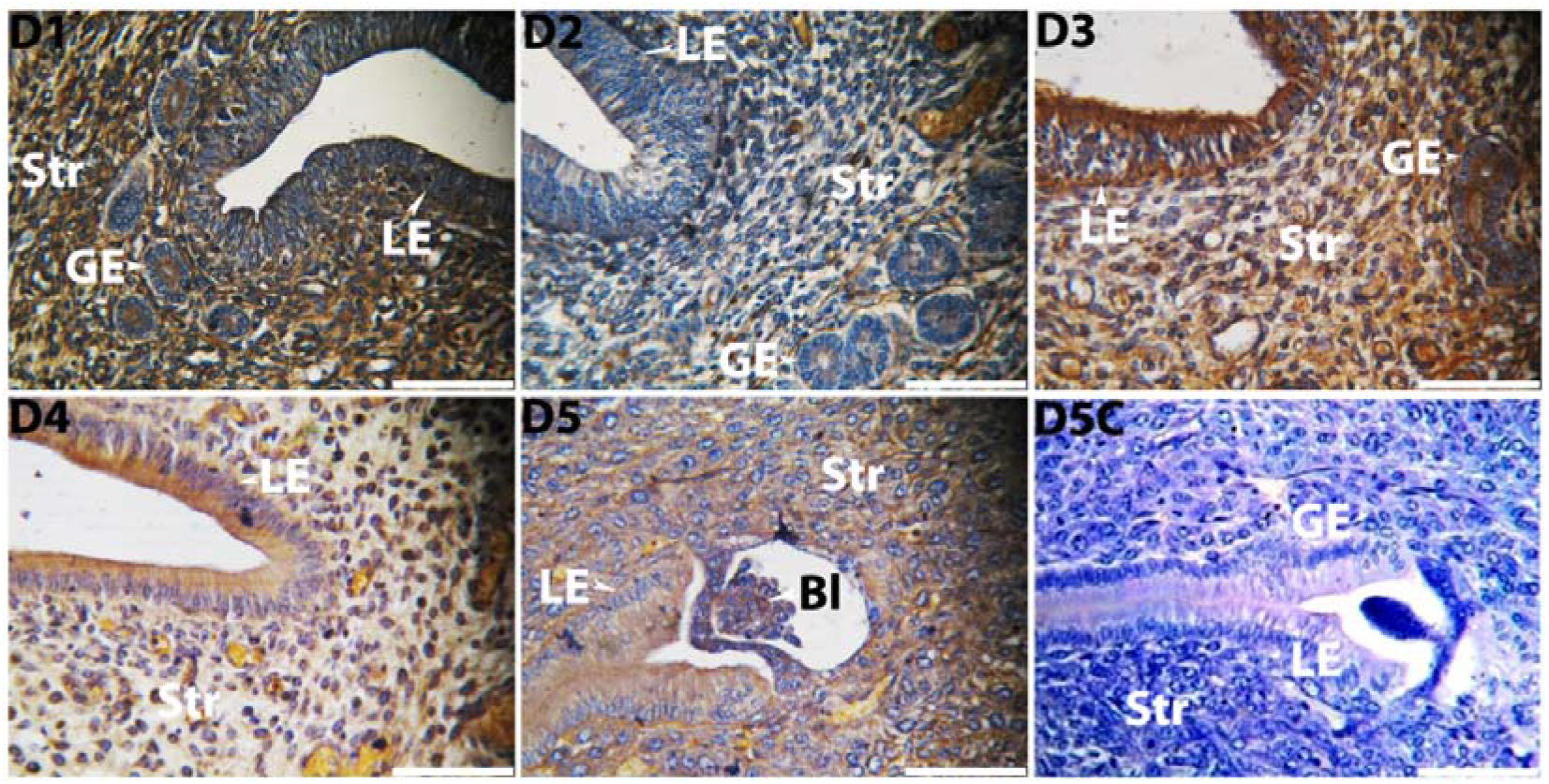
Immunohistochemical localization of cytochrome P_450_ aromatase protein in the uterus of golden hamster during pre-implantation (D1 to D3) and peri-implantation (D4 and D5) period showing their spatiotemporal localization in stroma (Str), luminal epithelium (LE) and glandular epithelium (GE). Brown deposition represents positive staining. Primary antibody has been replaced by rabbit IgG in day 5 (D5) pregnant uterus and considered as negative control (D5C). Photographs were taken under bright field condition. Scale bar-50µm and n = 5 in all the groups. Bl represents blastocyst and D represent day of pregnancy. DM showing myometrium region of uterus in specified dated pregnancy and DC represent negative control of specific dated pregnancy.

**Fig. 3:**
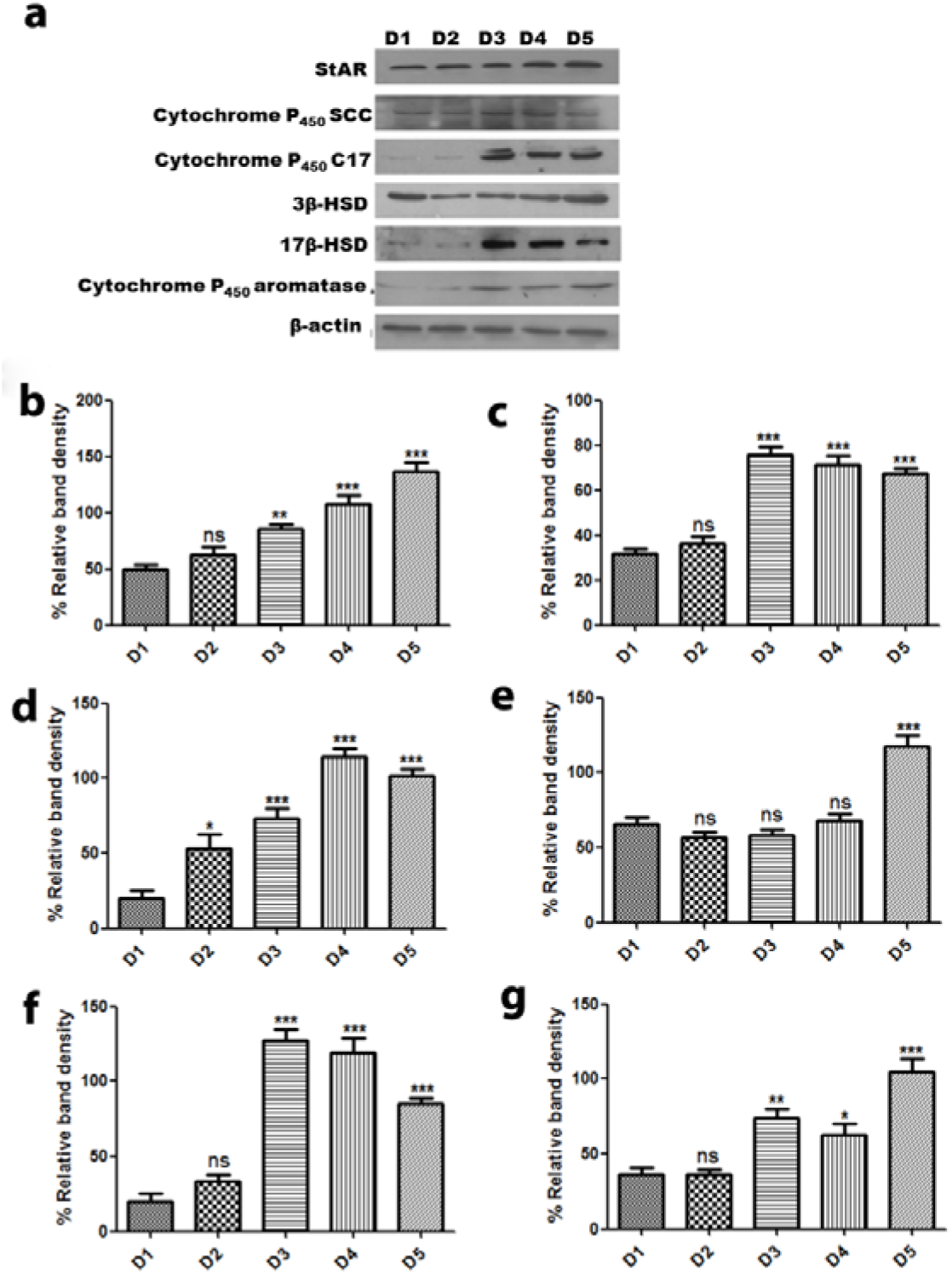
Western blot for enzymes involved in estrogen biosynthesis in uterus of golden hamster during pre and peri-implantation period. Fig.3 **a**. Represent blot of different enzymes involved in steroidogenic pathway in uterus. **Fig.3b. to** Fig.3 **g**. represents the densitometry analysis of blot for StAR, Cytochrome P_450_ SCC, Cytochrome P_450_ C17, 3β-HSD, 17β-HSD and Cytochrome P_450_ aromatase respectively. Graphs expressed as a percentage ratio of intensity of each enzyme to that of β-actin. All the data are expressed as mean ± SEM of three individual experiments. P value ≤ 0.05 is considered significant. (* for P ≤ 0.05, ** for P ≤ 0.01 and *** for P ≤ 0.001)

**Fig. 4:**
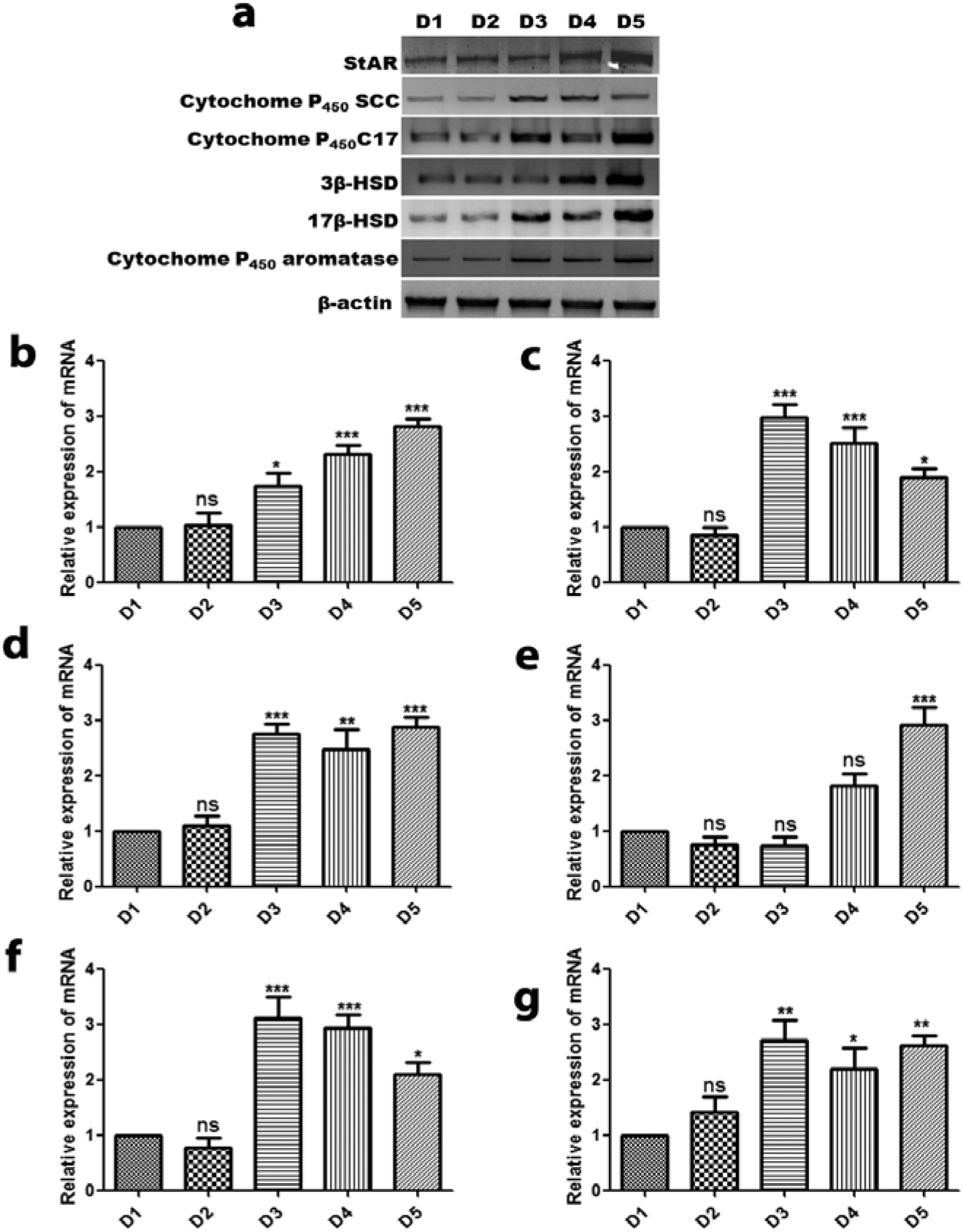
Quantitative and semi-quantitative PCR for mRNA of enzymes involved in estrogen biosynthesis in uterus of golden hamster during pre and peri-implantation period. **Fig.4a**. Represent gel image of semi-quantitative PCR of mRNA of different enzymes involved in steroidogenic pathway in uterus. **Fig.4b. to** Fig.4 **g**. represents the real time PCR analysis of mRNA for StAR, Cytochrome P_450_ SCC, Cytochrome P_450_ C17, 3β-HSD, 17β-HSD and Cytochrome P_450_ aromatase respectively. Fold changes are expressed in relative to day 1 of pregnancy. Data represent the mean ± SEM of three independent biological experiments. P value ≤ 0.05 is considered significant (* for P ≤ 0.05, ** for P ≤ 0.01 and *** for P ≤ 0.001)

More immunopositive cells of Cytochrome P_450_ scc were observed in day 3 of pregnancy as compare to other day of peri-implantation period. This enzyme was found to present only in luminal epithelia in D5 of pregnancy (Fig. 2B). Cytochrome P_450_ C17 was found to be present in the uterus during all the day of peri-implantation period with comparatively less immunopositive cells in D1 and D2 as compared to other day. It has more expression in D3 to D5 of pregnancy with maximum immunopositive cells in luminal epithelia of D5 of pregnancy (Fig. 2C).

Almost similar pattern of localization of 3β-HSD and 17β-HSD were observed in the uterus during peri-implantation period. They are found to be more localized in luminal epithelia, glandular epithelia and in stromal cells of D4 and D5 pregnant uterus (Fig. 2 D and E). Cytochrome P_450_ aromatase was also observed to localize in luminal epithelia in all the day of peri-implantation period with little expression in D2 of pregnancy (Fig. 2 F).

All these enzymes were also found to be present in uterus of D6 to D8 of pregnant hamster. In D6 uterus these were found to be present in whole stromal cells and luminal epithelia. However in D7 uterus it was observed only around the blastocyst. These enzymes were only present in trophectoderm in D8 of pregnancy with almost negligible expression in stromal cells (Fig. 5A and 5B).

**Fig. 5A:**
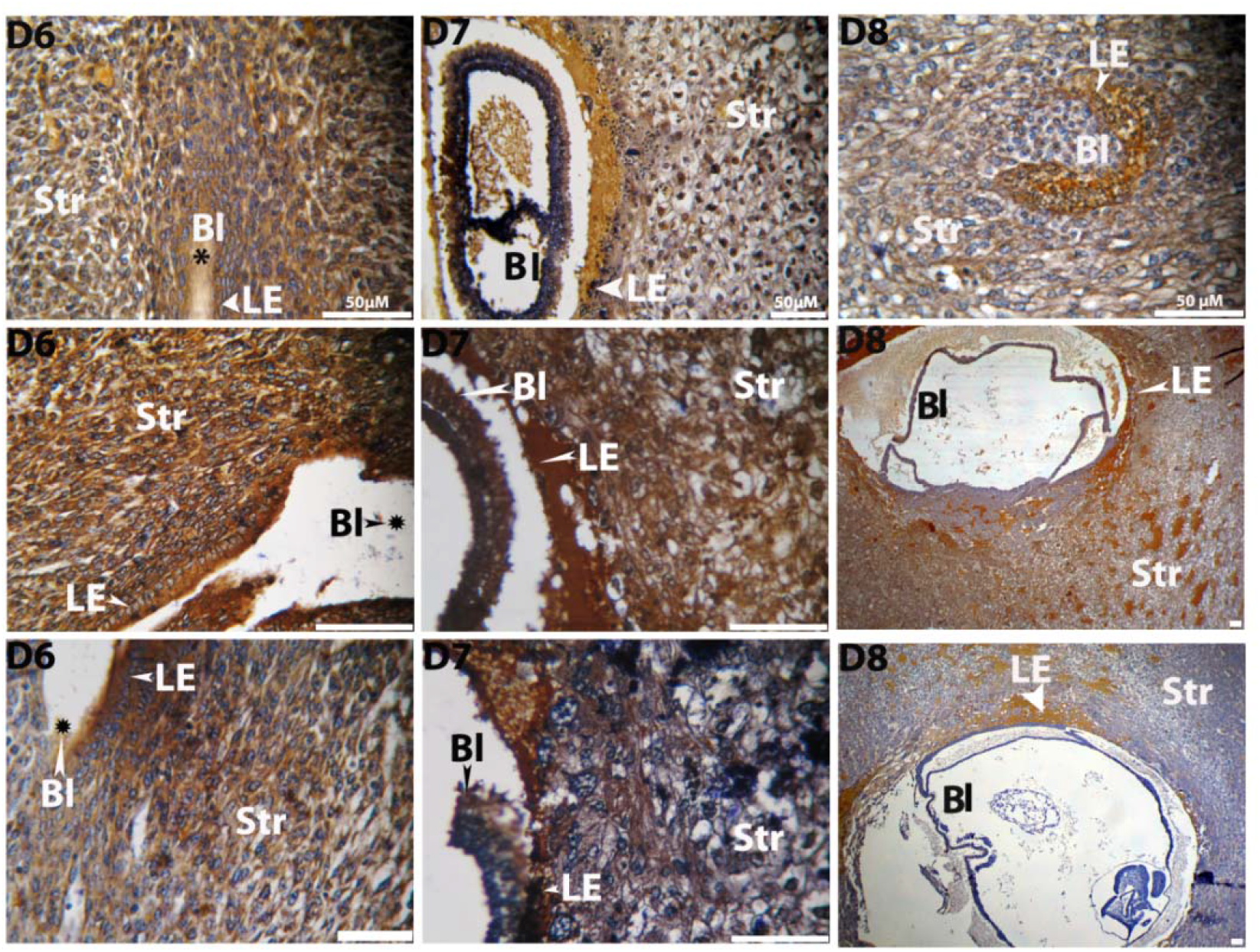
Immunohistochemical localization of StAR, cytochrome P_450_ scc and cytochrome P_450_ C17 protein respectively in the uterus of golden hamster during post-implantation (D6 to D8) period showing their spatiotemporal localization in stroma (Str), luminal epithelium (LE) and glandular epithelium (GE). Brown deposition represents positive staining. Photographs were taken under bright field condition. Scale bar-50µm and n = 5 in all the groups. Bl represents blastocyst and D represent day of pregnancy.

**Fig. 5B:**
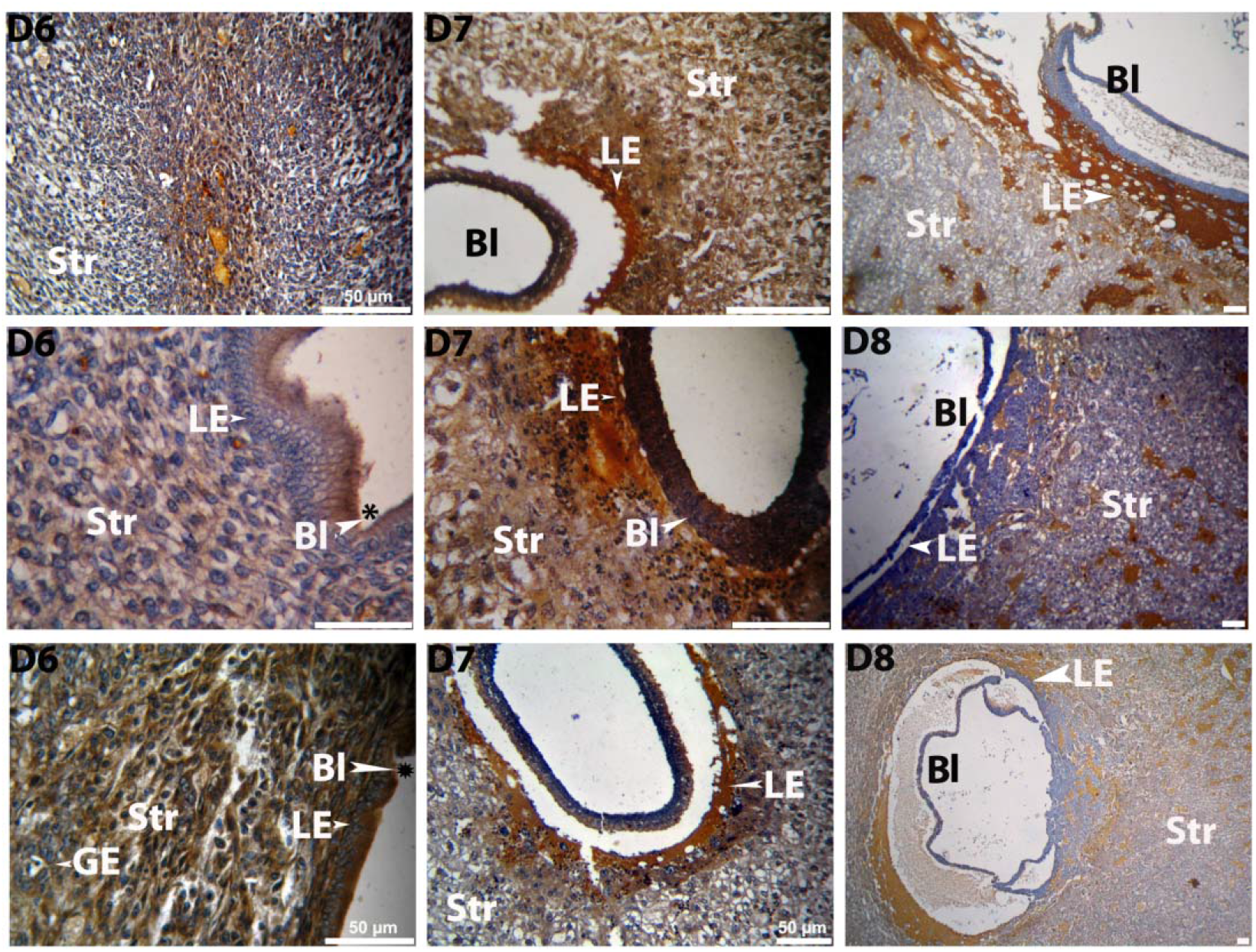
Immunohistochemical localization of 3β-HSD, 17-β HSD and cytochrome P_450_ aromatase proteins respectively in the uterus of golden hamster during post-implantation (D6 to D8) period showing their spatiotemporal localization in stroma (Str), luminal epithelium (LE) and glandular epithelium (GE). Brown deposition represents positive staining. Photographs were taken under bright field condition. Scale bar-50µm and n = 5 in all the groups. Bl represents blastocyst and D represent day of pregnancy.

### Expression of different enzymes and their mRNA of estrogen biosynthesis pathway in early pregnant uterus

These enzymes were found to be expressed in all the day of early pregnancy. Their expression was found to be significantly upregulated (P ≤ 0.05) around the day of implantation (D3 to D5) as compared to D1 of pregnancy indicating their role to support the process of embryo implantation (Fig.3). Their expression is also found to be upregulated in D6 of pregnancy indicating their positive contribution in the process of decidualization (Fig. 6).

**Fig. 6:**
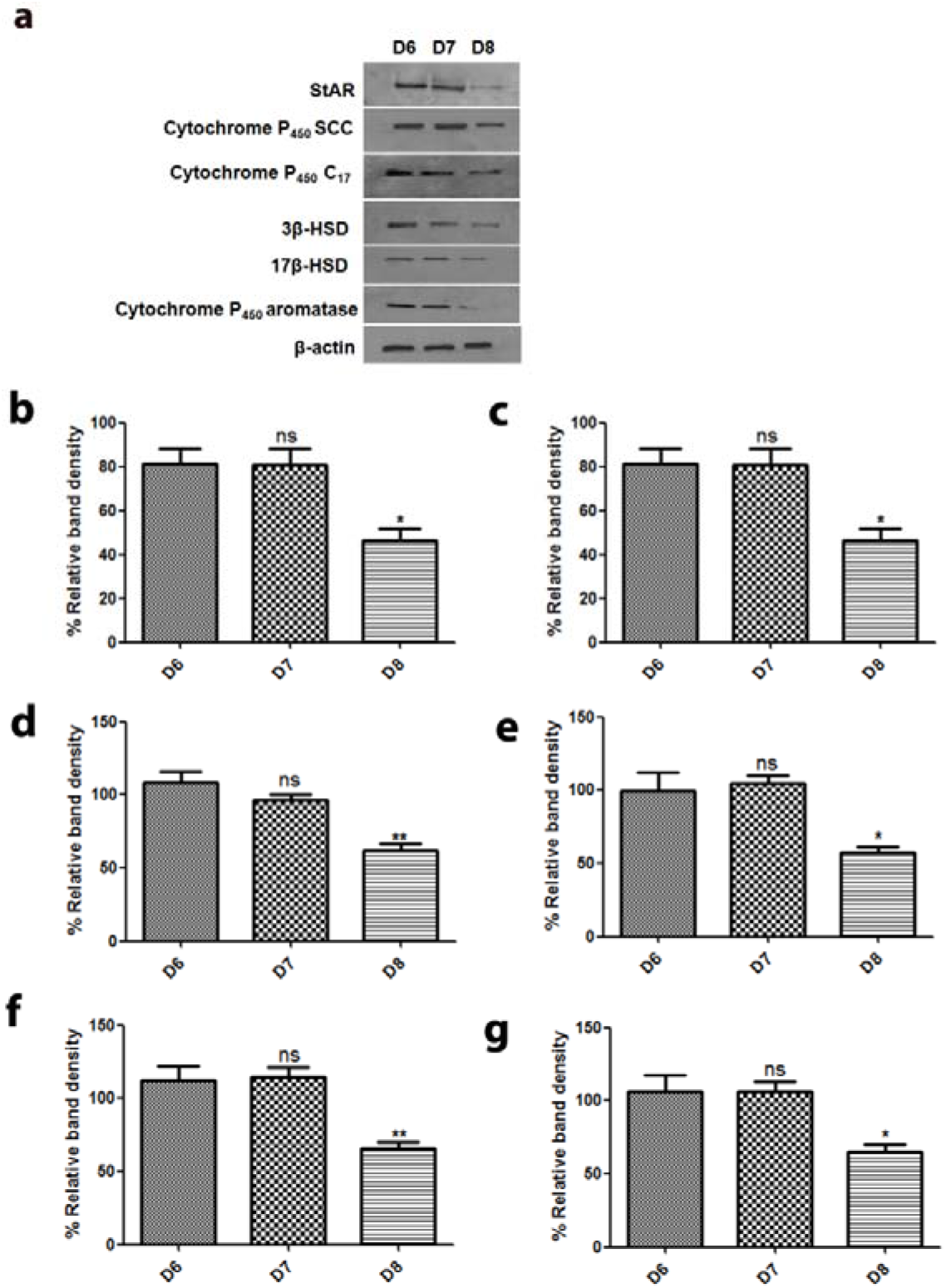
Western blot for enzymes involved in estrogen biosynthesis in uterus of golden hamster during post-implantation period. Fig.6 **a**. Represent blot of different enzymes involved in steroidogenic pathway in uterus. **Fig.6b. to Fig.6g**. represents the densitometry analysis of blot for StAR, Cytochrome P_450_ SCC, Cytochrome P_450_ C17, 3β-HSD, 17β-HSD and Cytochrome P_450_ aromatase respectively. Graphs expressed as a percentage ratio of intensity of each enzyme to that of β-actin. All the data are expressed as mean ± SEM of three individual experiments. P value ≤ 0.05 is considered significant (*for P ≤ 0.05, ** for P ≤ 0.01 and *** for P ≤ 0.001).

Their mRNA expression profile corroborates the finding in western blot (Fig. 4 and Fig. 7). They are found to be upregulated around the day of implantation and decidualization.

**Fig. 7:**
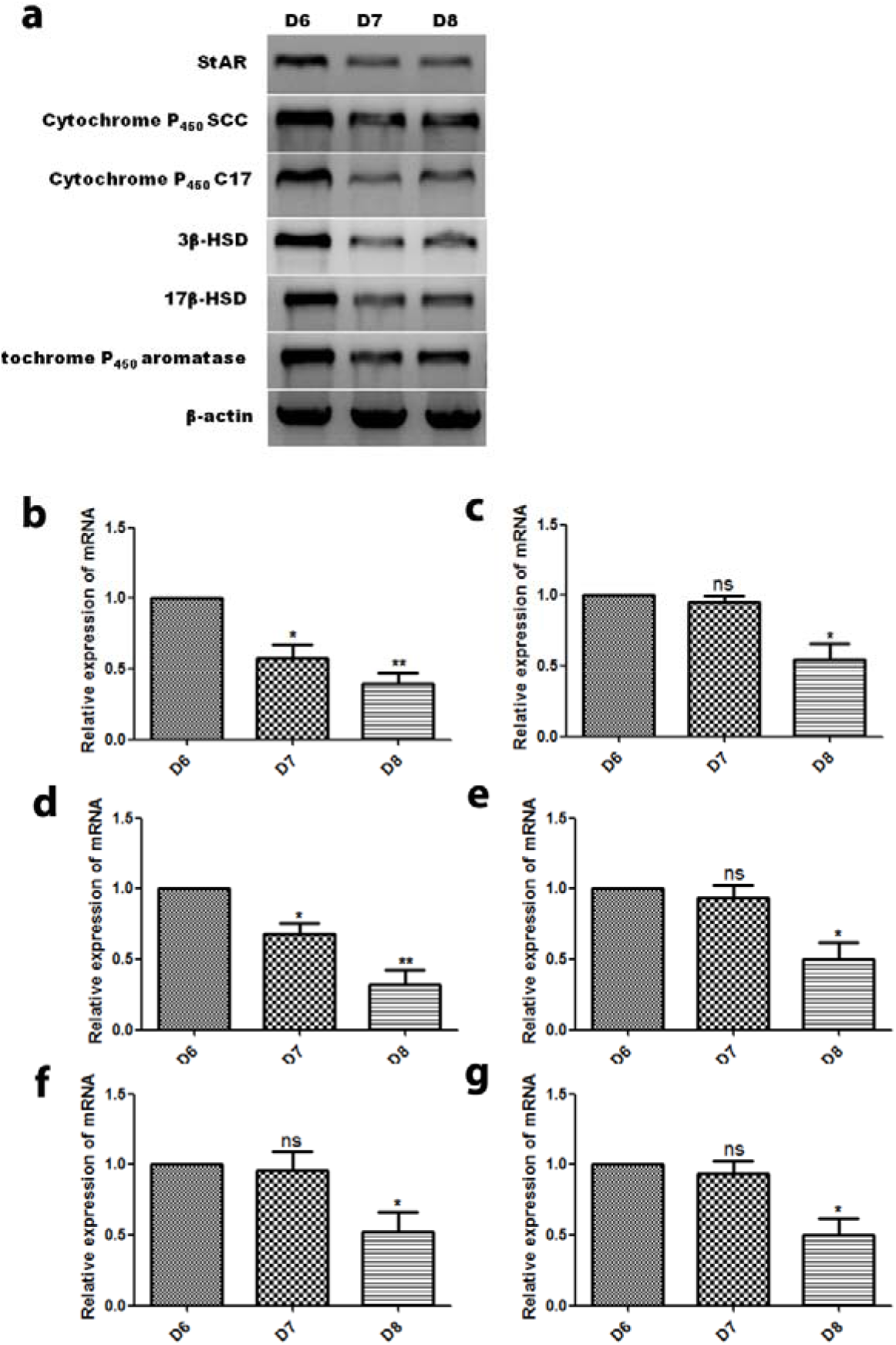
Quantitative and semi-quantitative PCR for mRNA of enzymes involved in estrogen biosynthesis in uterus of golden hamster during post-implantation period. **Fig.7a**. Represent gel image of semi-quantitative PCR of mRNA of different enzymes involved in steroidogenic pathway in uterus. **Fig.7b. to** Fig.7 **g**. represents the real time PCR analysis of mRNA for StAR, Cytochrome P_450_ SCC, Cytochrome P_450_ C17, 3β-HSD, 17β-HSD and Cytochrome P_450_ aromatase respectively Fold changes are expressed in relative to day 6 of pregnancy. Data represent the mean ± SEM of three independent biological experiments. P value ≤ 0.05 is considered significant (*for P ≤ 0.05, ** for P ≤ 0.01 and *** for P ≤ 0.001).

## Discussion

Embryo implantation is the most critical step for the establishment of pregnancy and considered as a most vulnerable point for the loss of early pregnancy. The event of embryo implantation is precisely orchestrated that occurs within a defied time frame and strictly regulated by estrogen and progesterone (Yang et al., 2016). Receptivity of endometrium is considered as crucial determinant for successful implantation which is regulated by ovarian estrogen and progesterone in mice (Wang et al., 2006). Ovarian estrogen and progesterone both are essential for the success of this process (Paria et al., 1993; McCormack et al., 1974 and Psychoyos et al., 1961). Early report of experimentally induced delay implantation model reflects the role of estrogen in mice. Ovariectomized day 4 pregnant mice were failed to implant embryo even after continual supply of progesterone while a single dose of external estrogen was found to be sufficient to initiate the process of embryo implantation (Yoshinga et al., 1966). But in majority of species including golden hamster ovarian estrogen are not found to be essential for implantation (Orsini et al., 1962; Harper et al., 1969; Greenwald et al., 1985; Deanesly et al., 1960; George et al., 1978 and Perry et al., 1973). However the process of delayed implantation is not observed in hamster as in mice. Absence of delayed implantation in golden hamster indicates either estrogen independent process of implantation or the involvement of estrogen from different source other than ovary. Simultaneously, adverse effect of estrogen receptor antagonist treatment on the embryo implantation of golden hamster (Sengupta et al., 1983) reflects a obligatory role of estrogen in this process. Our previous result of intrauterine treatment of ICI-182780, an estrogen receptor antagonist, in day 3 of pregnancy completely inhibits this process in golden hamster (Kumar et al., 2017). Hence, there is possibility of involvement of non-systemic estrogen for the establishment of early pregnancy in golden hamster. In this study, we have considered pregnant uterus as a novel site for *de novo* biosynthesis of estrogen which is sufficient to induce estrogen responsive physiological function to initiate the event of early pregnancy. Further, to confirm the biosynthesis of uterine estrogen and their positive role on embryo implantation, we administered letrozole, a inhibitor of cytochrome P_450_ aromatase, a rate limiting enzyme for the synthesis of estrogen, intrauterine just prior to implantation and found the complete absence of implantation in treated horn while vehicle treated control uterine horn have healthy implantation site. Our observation is in line with to the finding of Das et al. in mice where they observed differed decidualization when aromatase inhibitor was injected intraluminally just after embryo implantation. This suggests the possibility of synthesis of estrogen in uterus of golden hamster during preimplantation stage which is essential for priming uterus for successful embryo implantation. When preimplantation uterus is treated with an aromatase inhibitor, it significantly checked the synthesis of estrogen that leads to failure of implantation in golden hamster.

The activity of several mitochondrial enzymes is essential for de novo synthesis of steroid hormones. Cytochrome P455scc is considered as a key steroidogenic enzymes that are responsible for the conversion of cholesterol to pregnenolone (Hall et al., 1984 and Miller et al., 1988). Our immunohistochemical localization suggests the presence of this enzyme in the early pregnant uterus of golden hamster in a spatiotemporal manner. Our results is consistent with the previous available reports in mice, rat as well as in human placenta (Schiff et al., 1993; Ben-Zimra et al., 2002 and Jonathan et al., 1999) where the activity of these enzymes were observed and supposed to be involved in the synthesis of progesterone during pregnancy. Later on, Das et al. showed that this enzyme in uterus is also associated with the biosynthesis of estrogen during the process of decidualization (Das et al., 2009). We also observed that the expression of this enzyme and their mRNA is upregulated in the uterus around the day of implantation and decidualization. It indicates that spatiotemporal localization and expression of this enzyme in the uterus of pregnant golden hamster is associated with the biosynthesis of estrogen. The efficiency of steroid hormone production depends on the supply of the substrate molecules, cholesterol, from the outer mitochondrial membrane to inner membrane where the key enzyme of this pathway resides. This transport of cholesterol from outer membrane is regulated by a protein commonly known as steroidogenic acute regulatory protein (StAR) encoded by a gene STARD1 (Stocco et al., 1996 and Clark et al., 1994). The expression of StAR is supposed to rapidly induce by steroidogenic stimuli and then it catalyze the intermembrane transfer of cholesterol to P450 scc and in this manner it begin steroid biosynthesis (Holly et al., 2009). STARD1 expression is not observed in human placenta however it has tremendous capacity of steroid biosynthesis at the end of pregnancy reflecting from very high level of progesterone (250 mg/day) at the end of pregnancy (Jaffe et al., 1999). It has been suggested that placental steroidgenesis is supported by a homologue of STARD1 known as MLN64 which function in similar way as like StAR protein (Tuckey et al., 2009 and Watari et al., 1997). C-terminal domain of MLN64 contain StAR domain which is homologous to StAR protein (Hill et al., 2014 and Folligan et al., 2011). MLN64, a placental constitutive protein containing a C-terminal START domain homologous to StAR protein and a number of other lipid transfer protein. We also observed the presence of StAR protein in the uterus of golden hamster during the process of embryo implantation and decidualization. The induced expression of StAR protein and their mRNA in the uterus during early pregnancy suggested their role in uterine estrogen biosynthesis. Second step of steroidogenic pathway is the formation of progesterone from pregnelone by the activity of 3β-hydroxysteroid dehydrogenase. This is a membrane bound enzmes and their enzymatic activity has been located in microsomes and mitochondrial fraction (Tuckey et al., 2005 and More et al., 2013). It has been assumed that endoplasmic reticulum fraction of this enzyme is responsible for the conversion of pregnenolone in to progesterone while mitochondrial faction catalyze the synthesis of androstenodione from DHEA (Yves et al., 2016). There are multiple isoform of 3β-HSD in human and rodents. All isoforms are encoded by a distinct gene Different isoforms are observed to express in tissue and time specific manner (Peng et al., 2012). 3β-HSD type VI is assumed to express in uterus of murine and found to be present at implantation site in the uterus of mice during early pregnancy (Abbaszade et al., 1997). Their expression was observed to increase during day 10 to day 12 of pregnancy in rodents (Durkee et al., 1992). We also demonstrate the presence of 3β-HSD in the uterus of golden hamster during peri and post-implantation period suggesting their role as a mediator in the steroid biosynthesis. There expression was found to be upregulated during window of implantation as well as post implantation period. This strengthens our hypothesis of uterine estrogen synthesis and their crucial role in the embryo implantation and decidualization. We also observed the localization and expression of 17β-HSD in spatiotemporal manner during early pregnancy in golden hamster. This enzyme was found to be essential for the conversion of androstenedione and testosterone as well as estrone and estradiole (Pepe et al., 2008). Four different isoforms of this enzyme has been observed and found that 17β-HSD-1 and 17β-HSD-2 are expressed in the placenta of human being. Our results also suggest the expression of this enzyme in the uterus during pregnancy. The subsequent conversion of 17β-HSD catalyzed product in to estrogen is achieved by the activity of an enzyme cytochrome P_450_ aromatase. This is a rate limiting enzyme in the biosynthesis of estrogen. This enzyme is responsible for the formation of phenolic ring through a series of reaction from C19 steroids substrate. Pregnant uterus of mouse was observed to express this enzyme during the process of decidualization. It was established that cytochrome P_450_ aromatase driven biosynthesis of estrogen in the uterus is responsible for decidualization (Das et al., 2009). We also observed the expression of cytochrome P_450_ aromatase in the uterus of golden hamster and found upregulated expression of this enzyme during early pregnancy. Together we can say that uterus of golden hamster expressed all the set of enzyme which involves in de novo synthesis of estrogen during early pregnancy. Expression of all these enzymes was found to be upregulated around the day of implantation and decidualization. Therefore, these data support our hypothesis that de novo synthesized estrogen in the uterus of golden hamster during early pregnancy is sufficient to elicit the physiological response required for the process of embryo implantation and decidualization.

